# Prediction of Inefficient BCI Users based on Cognitive Skills and Personality Traits

**DOI:** 10.1101/2021.09.28.461955

**Authors:** Laura J. Hagedorn, Nikki Leeuwis, Maryam Alimardani

## Abstract

BCI inefficiency is one of the major challenges of motor imagery brain-computer interfaces (MI-BCI). Past research suggests that certain cognitive skills and personality traits correlate with MI-BCI real-time performance. Other studies have examined sensorimotor rhythm changes (also known as *μ* suppression) as a valuable indicator of successful execution of the MI task. This research aims to combine these insights by investigating whether cognitive factors and personality traits can make predictions of a user’s ability to modulate *μ* rhythms during a MI-BCI task. Data containing 55 subjects who completed a MI task was employed, and a stepwise linear regression model was implemented to select the most relevant features for *μ* suppression prediction. The most accurate model was based on these factors: Spatial Ability, Visuospatial Memory, Autonomy, and Vividness of Visual Imagery. Further correlation analyses showed that a novice user’s *μ* suppression during a MI-BCI task can be predicted based on their visuospatial memory ability, as measured by the Design Organization Test (DOT).

## 1 Introduction

Brain-Computer Interface (BCI) systems allow users to control external devices by decoding brain activity. A common approach to translate brain activity is through Motor-imagery Brain-Computer Interface (MI-BCI) where the user’s imagination of movement is mapped into the movement of the machine. MI-BCIs have been investigated in several applications ranging from communication and movement control to neuro-rehabilitation [6]. Therefore, MI-BCIs are especially relevant to researchers and practitioners in the field of healthcare, as they can support patients with distinct forms of motor impairment [7].

In contrast to their high potential, MI-BCIs are barely used outside laboratories. One obstacle in the implementation of MI-BCI systems is the inconsistency of user performances, with BCI inefficiency being one of the major challenges [16, 29, 2]. It has been shown that around 15% to 30% of subjects are not able to control MI-BCI systems even after extensive training, which can lead to frustrating and costly procedures for users as well as researchers [29]. Well-defined prediction paradigms that identify inefficient users early on in research could help to avoid unnecessary training sessions or lead to the establishment of alternative training protocols for the low performers [29]. Therefore, MI-BCI performance prediction is of high interest to the BCI community.

Several researchers have already investigated the impact of individual factors (e.g. gender, personality, cognitive ability, etc.) as well as temporal factors (e.g. motivation, fatigue, etc.) on MI-BCI performance [5, 10, 16, 18, 24, 32]. For instance, it has been established that users’ performance is influenced by mental and psychological states [10, 5, 8]. Jeunet et al. [5] found that tension and self-reliance strongly correlate with MI-BCI performance. Participants that scored a high value on the tension dimension were classified as highly tense, impatient, and easily frustrated personalities and showed a low MI-BCI performance. On the other hand, participants with high self-reliance and the ability to learn autonomously achieved a high BCI performance. Jeunet et al. [5] classified the learning style of participants, which requires a certain level of autonomy, as a crucial factor and reliable predictor of MI-BCI performance.

Additionally, many researchers have found correlations between cognitive profile of a user and their MI-BCI performance [5, 10, 18, 26]. For instance, Jeunet et al. [5] reported strong correlations between MI-BCI performances and user’s spatial ability as measured by a mental rotation task. Similarly, Pacheco et al. [18] found correlations between spatial visualization ability (as measured by Block Design Test and Mental Rotation Task) and performance of 7 male users in a flexion vs. extension MI-BCI.

More recently, Leeuwis et al. [9, 10] investigated the impact of spatial ability and visuospatial memory as well as personality traits on MI-BCI performance of 55 participants. While their experiment did not confirm the impact of spatial ability (measured by the Mental Rotation Task) on MI-BCI performance, they reported a significantly higher visuospatial memory (as measured by the Design Organization Task) in high aptitude BCI users. They further implemented a stepwise linear regression to select relevant predictors, and the best results were achieved by a model based on *Gender + Emotional Stability + Orderliness + Vividness of Visual Imagery*. They concluded that visuospatial ability correlates positively with the users’ performances, which was classified as a negative predictor by Jeunet et al. [5]. Because of these contradicting findings, spatial ability and visuospatial memory are reviewed in particular in this research.

The presented studies as well as other studies that aim to predict users’ MI-BCI performance mainly rely on the user’s online classification accuracy, which is the percentage of correctly recognized mental commands by a BCI classifier [12]. However, using this metric alone could be problematic as BCI classifiers are often calibrated at the beginning of the experiment, and then applied to classify the EEG signals in the following trials [12]. This means that the classifier parameters are not adapted to the subject’s learning of the MI task, and hence the low BCI accuracy could be an outcome of the system and not the user [1, 27].

To avoid this pitfall, it is suggested to rely on brain pattern changes in the sensorimotor cortex (in the *μ* band) that are associated with movement imagery as an indicator of MI performance [14]. It has been shown that the *μ* waves are larger when a subject is at rest and are suppressed when the person generates, observes or imagines movements [7, 20]. This pattern change is referred to as event-related (de)synchronization (ERD/ERs) (in short *μ* suppression) and is commonly used as a reliable measure in development of MI-BCI classifiers [4, 21, 31]. Based on this, our study focused on the following research question:

> *Can cognitive skills and personality traits predict a user’s ability to generate μ suppression in sensorimotor regions during motor imagery of left and right hands?*

To answer this question, we used a data set containing personality information, cognitive test results and EEG signals from 55 participants who completed a two-class motor imagery BCI experiment. Using EEG signals from C3 and C4, *μ* suppression values corresponding to the left and right-hand imagery were computed and together with personality and cognitive factors were used to train a stepwise linear regression model. This enabled selection of the relevant factors that could best predict a subject’s MI task performance.

## 2 Methods

### 2.1 Experiment

#### 2.1.1. Participants

The data employed in this research was collected by Leeuwis et al. [10]. EEG signals were collected form 55 participants (36*females, M_age_* = 20.71, *SD_age_* = 3.52) while they performed a two-class motor imagery task. The participants were all right-handed and naïve to MI-BCI, which means that they had no prior experience with either MI task or BCI systems.

#### 2.1.2. EEG recording

The EEG signals were recorded via 16 electrodes distributed over the sensorimotor area according to the international 10 – 20 system (F3, Fz, F4, FC1, FC5, FC2, FC6, C3, Cz, C4, CP1, CP5, CP2, CP6, T7, T8). The g.Nautilus amplifier was used to amplify the recorded signals, and the sampling rate was 250 samples per second. To minimize the noise during recording, a bandpass filter from 0.5 to 30 Hz was applied.

#### 2.1.3. Motor imagery task and the BCI system

Details of the experimental procedure can be found in Leeuwis et al. [10]. After participants completed the demographic questionnaire and the cognitive tests (Fig 1A), they were placed before a screen and the researcher positioned the EEG cap on their head (Fig 1B). The researcher explained that the participant had to imagine squeezing their left or right hand according to the cue on the screen without generating any physical movement or muscle tension. The BCI task was repeated in four runs, each including 40 motor imagery trials (20 left and 20 right). Each trial took 8 seconds starting with a fixation cross, followed by a cue at second 3, which indicated the direction of the MI task, and finally the presentation of feedback associated with the task during the last 3.75 seconds (Fig 1C). The first run was used for calibration and did not provide feedback. In runs 2 to 4, the parameters of the classifier were adjusted according to the EEG data collected in the latest run to provide feedback to the subject.

**Fig. 1.**
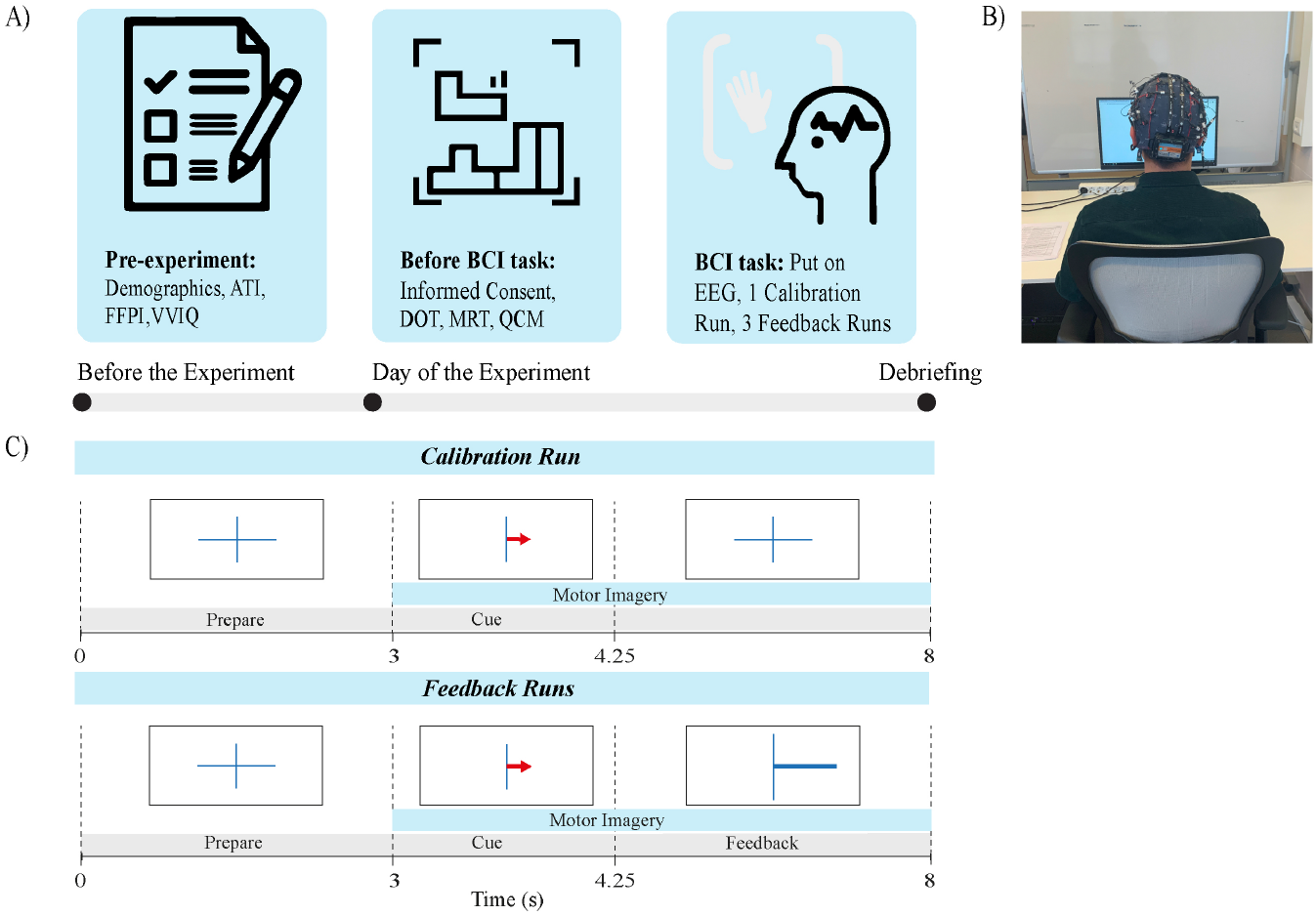
A) the procedure of the experiment, B) the experimental setup and C) the timeline of the MI task in one BCI trial. From [10].

#### 2.1.4. Measurements

Several individual factors including demographics, personality traits and cognitive skills were measured before the participants performed the BCI task (Fig 1A). These measures are summarized in Table 1. For a more detailed explanation and illustration of the survey procedure, see [10].

**Table 1.** Overview of the tests and questionnaires that were employed to measure certain cognitive skills and personality traits.

| Measurement | Description |
| --- | --- |
| Demographic questionnaire | Questionnaire to access background information such as age, gender, educational level, as well as participant's experience with video games, sports, and music. |
| Affinity for Technology Interaction Scale (ATI) | Proposed by [4] to evaluate an individual's tendency to actively engage with technology. |
| Mental Rotation Task (MRT) | A cognitive test to evaluate the subjects' spatial ability by measuring their ability to mentally rotate objects in space [6]. |
| Design Organization Test (DOT) | A cognitive test to measure the visuospatial memory of the participants [10]. |
| Five Factor Personality Inventory (FFPI) | A questionnaire to assess the Big Five factors of an individual's personality. These include Extraversion, Autonomy, Orderliness, Emotional stability, and Mildness [7]. |
| Vividness of Visual Imagery Quiz (VVIQ) | A psychometric assessment, proposed by [17], to measure a subject's mental imagery skills. Participants are asked to conceptualize certain people, objects, or settings in their minds. |
| Questionnaire for Current Motivation (QCM) | An assessment to measure an individual's motivation to learn and use BCI systems. |

### 2.2 Data Analysis

Data analysis was conducted in Python (version 3.9.1). This included the EEG pre-processing, *μ* suppression analysis and the implementation of the machine learning algorithm.

#### 2.2.1. EEG pre-processing and *μ* suppression analysis

Similar to [11] and [19], *μ* rhythms were exclusively obtained from the EEG channels C3 and C4, as the *μ* suppression is the strongest in these areas [11, 23]. Given that *μ* waves are constricted to the movement of the opposite upper limb, the brain activity at C4 was associated with the left-hand MI and activity at C3 with the right-hand MI.

Once the signals from these channels were isolated, they were segmented into MI trials. In each trial, the first 3 seconds was extracted as the resting state, and the last 4 seconds was identified as the MI phase (Fig 1C). For each EEG segment (rest and MI), spectral density was estimated using Welch’s periodogram [13] and the mean power over the *μ* frequency band (8-12Hz) was computed. Thereafter, the event-related (de-)synchronization for each trial was calculated using the following function [19]:

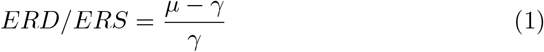

where *μ* represents the average power during the MI phase and *γ* is the average power during the rest period. This procedure was repeated for all participants separately for the left-hand (C4) and right-hand (C3) MI trials in all 4 runs. Finally, the resulted *μ* suppression for each participant in all trials was averaged and assigned as the participant’s final *μ* suppression value. Once the final *μ* suppression values for all participants were calculated, outliers were detected by using the interquartile range technique, which detected and removed two outliers (subject 13 and 14) in the data.

#### 2.2.3. Regression model

Because of the large number of predictors that were investigated in this research, a stepwise linear regression algorithm was implemented. This model iterates through the predictor variables and only includes features that improve the model. Additionally, it repeatedly controls whether the significance level of earlier added features decreases and removes them through backward elimination if that is the case [25]. In the present research, the maximum number of features to be selected at one time was set to four. The subsets of features were evaluated on their coefficient of determination (*R*^2^), which is a commonly used measurement for the accuracy of predictive regression models, as it measures how well the predictions of a model fit the actual values [15]. The *R*^2^ value ranges between 0 and 1, with 1 referring to a model that explains all the variation in the given predictor variables.

#### 2.2.4. Correlation analysis

Following the identification of the most relevant features by the stepwise linear regression model, a correlation analysis was performed on the selected features separately. Pearson’s r was calculated for normally distributed variables and Kendall’s tau for variables with non-normal distribution. Differences of gender were checked with an independent t-test. The variables were checked for normality by applying the Shapiro-Wilk test [22]. Multicollinearity was tested with the Variance Inflation Factor (VIF) similar to Leeuwis et al. [10].

## 3 Results

Our results showed no significant differences between genders (*t*(34) = –0.19, *p* = 0.8) in terms of generated *μ* suppression, and neither did the age of the participants (*r_τ_* = 0.028, *p* = 0.7).

The model that was able to predict *μ* suppression the most accurately was generated by: *Spatial Ability + Visuospatial Memory + Autonomy + VVIQ* (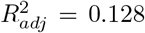, *p* < .05). Further correlation tests on the identified features in this model showed that Visuospatial Memory (*r*(50) = .32, *p* = .02) significantly correlated with *μ* suppression (Fig. 2). However, no significant correlation was found between Spatial Ability (as measured by the MRT) (*r*(50) = .008, *p* > .5), Autonomy (*r*(50) = 0.09, *p* < .5), VVIQ (*r*(50) = – .1, *p* < .5) and the generated *μ* suppression.

**Fig. 2.**
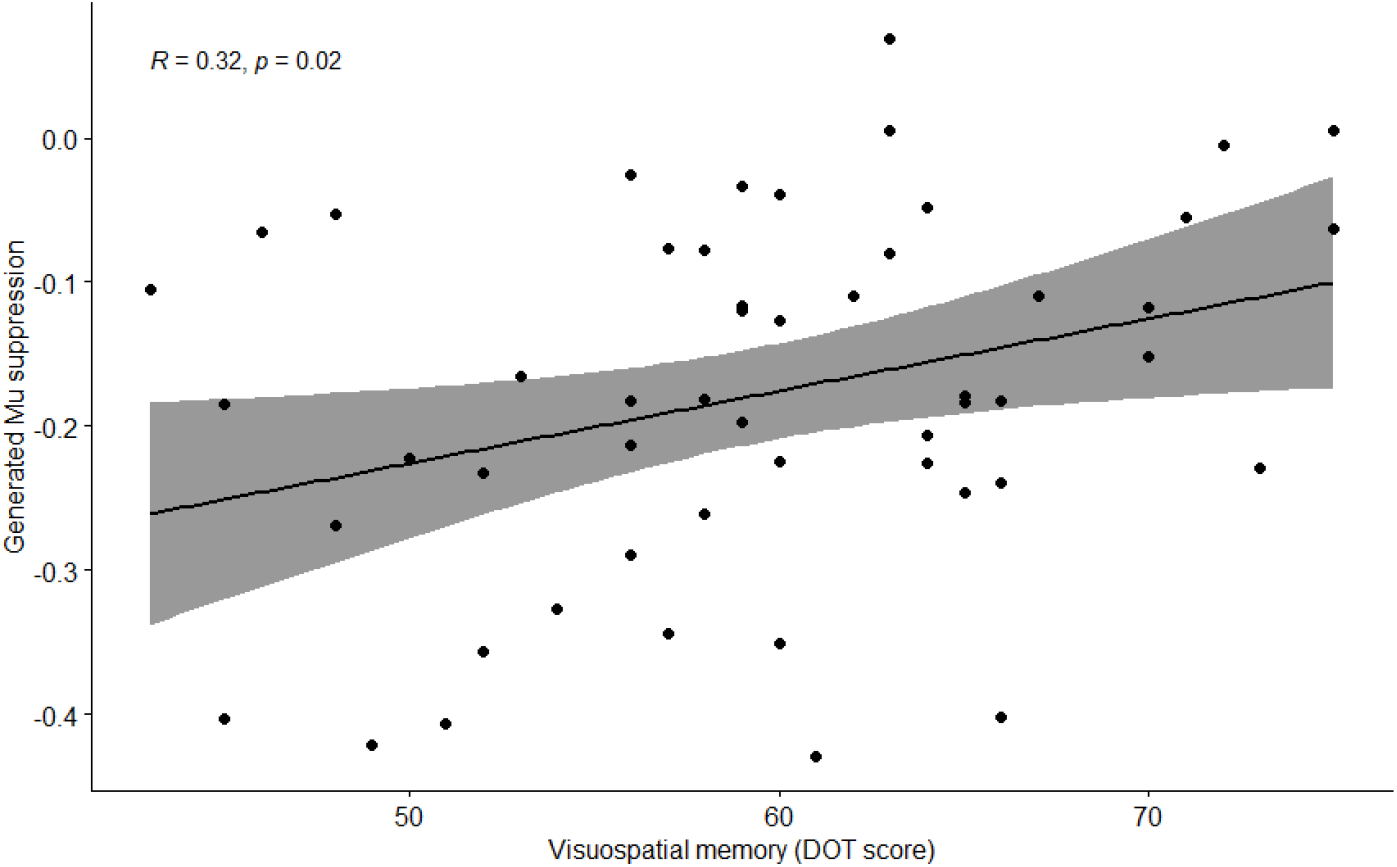
Significant correlation was found between participants’ visuospatial memory (DOT score) and *μ* suppression value in a MI-BCI task.

Finally, the best performing model proposed by Leeuwis et al. [10] (Gender + Emotional Stability + Orderliness + VVIQ) was tested, but did not show significant results in terms of prediction of *μ* suppression (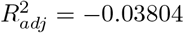, *p* = 0.7).

## 4 Discussion

The aim of this research was to investigate whether the cognitive skills and personality traits can be used as effective predictors of MI-BCI (in)efficiency among novice BCI users. Unlike previous studies that relied on the BCI classifier’s online performance as a measure of the user’s efficiency score [5, 10], we used the subject’s ability to produce *μ* suppression as an indicator of their MI task performance [19].

The *μ* band ERD/ERS of EEG signals at C3 and C4 during a left vs. right MI-BCI task were extracted from 55 participants, and after outlier removal, a stepwise linear regression algorithm was implemented to predict 53 participants’ *μ* suppression value based on their cognitive skills and personality traits. The best performing model obtained by our study identified four factors of Spatial Ability, Visuospatial Memory, Autonomy and Vividness of Visual Imagery as predictors of *μ* suppression. Among these, the only cognitive factor that demonstrated a significant correlation with *μ* suppression value was Visuospatial Memory. Although, the rest of features did not correlate with the generated *μ* suppression individually, their inclusion in the model supports the finding of previous research [5, 10].

Leeuwis et al. [9], as well as Pacheco et al. [18], found that high aptitude subjects performed better on the Design Organization Test that was used to measure visuospatial ability. The current research found that visuospatial memory (as measured by the DOT) correlates negatively with the magnitude of *μ* suppression, which means that subjects that scored higher on the DOT generated more *μ* suppression compared to subjects with lower scores on the DOT. Similar outcomes were detected by Jeunet et al. [5] who used the Corsi Block task to evaluate visuospatial short-term and working memory abilities and classified visuospatial memory as a negative predictor of MI-BCI performance. Further research is necessary to ascertain the role of visuospatial memory skills on MI and BCI performance and resolve the inconsistency that exists in these reports.

Additionally, past research reported relationships between spatial ability as measured by the Mental Rotation Test (MRT) and MI-BCI performance [5, 18]. Although no prior study has investigated the *μ* rhythms with regard to spatial ability and BCI performance, it was expected that participants who scored higher on the MRT would also show a higher ability in generating *μ* suppression. This assumption was based on the establishment that spatial ability is intimately related to the concept of motor imagery [5, 17]. However, this research did not validate this hypothesis, as there was no significant correlation between MRT scores and *μ* suppression.

Moreover, Jeunet et al. [5] concluded that MI-BCI performance is correlated with the users’ ability to learn independently, which requires a certain level of autonomy. Leeuwis et al. [10] further confirmed personality dimension Autonomy as an effective predictor of MI-BCI performance. Although Autonomy, as an independent factor, was not significantly correlated with *μ* suppression, it was chosen as a relevant feature by the regression model, indicating that it significantly improved the performance of the model and can therefore be considered as a potential predictor for MI-BCI performance in the future research.

In the experiment conducted by Leeuwis et al. [10], high vividness of visual imagery was a positive predictor of MI-BCI performance. Similarly, Touckovic and Osuagwu [30] reported a positive correlation between MI-BCI performance and vividness of imagery. The results of this study partially supported the impact of this factor on MI performance by reporting VVIQ as a predictor in the best performing model. However, further correlation analysis did not show significant relationship between VVIQ and *μ* suppression. This result is consistent with the reports of Vasilyev et al. [28] who also did not find a correlation between vividness of imagery and *μ*-rhythm ERD, but observed a significant impact of kinaesthetically vivid images on subjects’ cortical excitability. Future research should further investigate the role of this factor, as these reports remain inconclusive.

Leeuwis et al. [10] proposed a linear regression model to predict users’ MI-BCI performance using the classifier’s online error rates. Their most accurate model was based on Gender + Emotional Stability + Orderliness + VVIQ. Although this research made use of the same data and implemented a similar stepwise linear regression model, their best performing model did not show significant prediction in the present study. The only factor that was common between their model and the best performing model in our study was Vividness of Visual Imagery (VVIQ). Gender, Emotional stability, and Orderliness did not correlate with *μ* suppression independently. This inconsistent finding calls for further research to fully understand the occurrence of the *μ* rhythms and its relationship with cognitive factors and MI-BCI performance.

Finally, some limitations of this study have to be mentioned. Although the data that was employed by this study included a larger number of subjects than other similar studies [5, 16], the data was imbalanced in terms of age and gender, which occurred due to convenience sampling [10]. Only young and healthy subjects participated in the study, which reduces the generalizability of our findings to other populations for which MI-BCI applications are more relevant (e.g. individuals with motor impairment). Although, studies have shown no difference in MI-BCI training between healthy subjects and patients with motor disabilities [3], the outcome of our study should be validated with other populations, particularly the motor impaired subjects. Future research should also focus on other cognitive profiles and psychological measures across a broad range of subjects in order to investigate the behaviour of the *μ* waves in relation to motor imagery and BCI performance. This will encourage development of accurate models that can predict a user’s (in)ability to operate a MI-BCI system as a step toward alleviation of the BCI inefficiency problem. Consequently, this will broaden the range of MI-BCI applications outside the laboratories and will support the implementation of reliable BCIs in clinical systems.

## 5 Conclusion

This study aimed to identify cognitive skills and personality traits that could predict a BCI user’s ability to generate *μ* suppression during a left vs. right-hand motor imagery task. Fifty-five subjects participated in a MI-BCI experiment in which their personality traits and several cognitive abilities were measured. A stepwise linear regression model was trained on the data to exclusively select relevant factors that could predict participants’ *μ* suppression associated with the MI task. Results showed that the most accurate model was based on three cognitive factors of Spatial Ability, Visuospatial Memory and Vividness of Visual Imagery and one personality trait, i.e. Autonomy. Subsequently, correlation analyses were performed for each of these predictors, resulting in a significant correlation between Visuospatial Memory and *μ* suppression. Our findings confirm that an individual’s personality and cognitive abilities can indeed predict their performance on the MI task. The outcome of this study will contribute to future MI-BCI research by providing a tool for identifying low BCI performers in an early stage of research or clinical treatment. This would ultimately save resources and facilitate adjusted user training for different groups of BCI users.

